# Perinatal features of Prader-Willi syndrome: a Chinese cohort

**DOI:** 10.1101/568451

**Authors:** Lili Yang, Qiong Zhou, Bo Ma, Shujiong Mao, Yanli Dai, Mingqiang Zhu, Chaochun Zou

## Abstract

**Background:** Prader-Willi syndrome (PWS) is a rare complex genetic disorder caused by an absence of expression of imprinted genes on the paternally derived chromosome 15q11-q13 region. This study aimed to characterize the perinatal features in a cohort of Chinese individuals with PWS.

**Methods:** We analyzed anonymous data of 134 patients from the PWS Registry in China. Perinatal and neonatal presentations were analyzed, and compared between the two PWS genetic subtypes. We also compared the perinatal features of PWS patients with the general population and other previous reported large cohorts from France, UK and USA.

**Results:** This study included 134 patients with PWS (115 patients with 15q11-q13 deletion and 19 with maternal UPD). High mean maternal age was found in this cohort (30.5 vs. 26.7) comparing with the general population. 88.6% of mothers reported a decrease of fetal movements. 42.5% and 18.7% of mothers had polyhydramnios and oligohydramnios during pregnancy, respectively. 82.8% of the patients were born by caesarean section. 32.1% of neonates had birth asphyxia, 98.5% had hypotonia and 97.8% had weak cry or even no cry at neonatal period. Feeding difficulty existed in 99.3% of the infants, and 94.8% of them had failure to thrive. 69.4% of the infants ever used feeding tube during hospitalization. However, 97.8% of them discontinued tube feeding after discharged home. Maternal age, maternal pre-pregnancy weight and BMI were significantly higher in the UPD group (all *P*<0.05).

**Conclusions:** PWS should be highlighted for differential diagnosis for infants with following perinatal factors including polyhydramnios, intrauterine decreased fetal movements, cesarean section, low birth weight, feeding difficulty, hypotonia, and failure to thrive. Higher maternal age may be a risk factor for PWS, especially for UPD, and further studies for the mechanism of PWS are required.

**Author Summary:** Early diagnosis and tailored multidisciplinary treatment are utmost important for better quality of life of the infants with Prader-willi syndrome. Genetic diagnosis for PWS is now easily available; and diagnosis can be made in most patients within the first months of life even prenatally if obstetricians or neonatologists can recognize the perinatal features of PWS well. However, most patients still had a late diagnosis because the early signs of PWS not recognized. Our study highlighted the perinatal features of a large cohort of Chinese patients with PWS, which will benefit for early diagnosis and treatment. We found high incidence of decreased fetal movements, polyhydramnios and delivery by cesarean section, and higher maternal age comparing with the general population. Neonatal features found in our cohort included low birth weight, birth asphyxia, failure to thrive, feeding difficulty, weak cry and hypotonia. We found mothers with UPD had higher maternal age, pre-pregnancy weight and BMI. Most patients required tube feeding during hospitalization, however tube feeding was discontinued by their parents after discharge at home. Nutrition deficiency was a serious problem in infants with PWS. Home tube-feeding instruction should be carried out for parents of PWS patients in China.

## Introduction

Prader-Willi syndrome (PWS) is a rare complex genetic disorder characterized by severe hypotonia and feeding difficulties in early infancy, followed excessive eating and gradual development of morbid obesity in early childhood, together with a series of comorbidities, including short stature, typical facial dysmorphism, psychomotor delay, behavioral abnormalities and cognitive impairment [1]. It is caused by an absence of expression of imprinted genes on the paternally derived chromosome 15q11-q13 region. PWS has several genetic subtypes: deletion of the paternal copy of 15q11-13 in about 65% of the cases, maternal uniparental disomy (UPD) for chromosome 15 in about 30%, imprinting center defect in less than 5%, and very rare cases of translocation involving the chromosome 15q11-q13 region [2,3]. Genetic diagnosis for PWS is now easily available; and diagnosis can be made in most patients within the first months of life even prenatally if obstetricians or neonatologists can recognize the perinatal features of PWS well. Increased awareness among obstetricians and healthcare providers would allow earlier diagnosis and treatment of PWS by pediatricians/neonatologists [4]. Early diagnosis and tailored multidisciplinary treatment are utmost important for better quality of life of the infants with PWS as they ensure comprehensive advice to prevent obesity, and stimulation of cognitive and adaptive skills [4,5].

It is imperative to make more obstetricians, pediatricians/neonatologists be aware of the perinatal features of PWS. Our study aimed to characterize the perinatal features in a cohort of Chinese individuals with PWS. We also compared the perinatal features of PWS patients with the general population and other previous reported large cohorts from France, UK and USA. This is the first large study on patients with PWS in China.

## Results

This study included 134 patients with PWS including 115 (85.8%) with 15q11-q13 deletion and 19 with maternal UPD (14.2%). Among these patients, 73 (54.5%) were boys (62 with deletions and 11 with maternal UPD) and 61 (45.5%) girls (53 with deletions and 8 with maternal UPD). The mean age of diagnosis was 31.75 ± 4.72 months with a range from 10 days to 17 years.

The mothers of this cohort of patients had a higher maternal age (30.5 vs. 26.7) than general pregnant women in China. 87.9% (109/124) of mothers of the patients reported a decrease of fetal movements during pregnancy. 42.5% and 18.7% of mothers had polyhydramnios and oligohydramnios during pregnancy, respectively. 82.8% of the patients were born by caesarean section, significantly higher than 34.9% in 2014 [7]. The rate of vaginal delivery was low of 17.2% (23/134), and 30% of them by forceps. 2.2% of mothers had hypertension during pregnancy, 5.2% had gestational diabetes, and 9.7% had premature rupture of membranes (Table 1).

**Table 1.** Perinatal factors of PWS and comparison with the general population in China

| Perinatal factors | Total | Population statistics |
| --- | --- | --- |
| Maternal variables |  |  |
| Maternal age, mean (range) | 30.5±5.5 (18-46) | 26.7±4.3<sup>[6]</sup> |
| Maternal pre-pregnancy BMI | 21.7±3.1 | 20.7±3.1<sup>[6]</sup> |
| Fetal movements (*n*=124): Decreased | 109 (87.9%) | NA |
| Normal | 14 (11.3%) | NA |
| Increased | 1 (0.8%) | NA |
| Pregnancy hypertension | 3 (2.2%) | 3.6%<sup>[6]</sup> |
| Gestational diabetes | 7 (5.2%) | 8.1%<sup>[6]</sup> |
| Preeclampsia | 0 (0%) | 0.52%<sup>[6]</sup> |
| Polyhydramnios | 57 (42.5%) | 1%-3%<sup>[6]</sup> |
| Oligohydramnios | 25 (18.7%) | 0.4-4.0%<sup>[6]</sup> |
| Premature rupture of membranes | 13 (9.7%) | 2%-4%<sup>[6]</sup> |
| Mode of delivery: Vaginal | 23 (17.2%) | 65.1%<sup>[7]</sup> |
| Caesarean section | 111 (82.8%) | 34.9%<sup>[7]</sup> |
| Neonatal variables |  |  |
| Gestational age: Preterm (< 37 weeks) | 22 (16.4%) | 7.1%<sup>[8]</sup> |
| Term (37- 41 6/7 weeks) | 102 (76.1%) |  |
| Post-term (≥42 weeks) | 10 (7.5%) | - |
| Birth weight: Normal | 85 (63.5%) | 93.8%<sup>[9]</sup> |
| Low birth weight | 44 (32.8%) | 7.2%<sup>[9]</sup> |
| Very low birth weight | 2 (1.5%) | - |
| Birth length: Boys | 48.5±3.2 | 46.9-54.0<sup>[10]</sup> |
| Girls | 49.4±3.7 | 46.4-53.2<sup>[10]</sup> |
| Birth asphyxia | 43 (32.1%) | 6.3%<sup>[6]</sup> |
| Hospitalization at neonatal period: Yes | 127 (94.8%) | NA |
| Duration, mean (range), day | 17 (0-90) | NA |
| Breast feeding | 15 (11.2%) | 58.5%<sup>[9]</sup> |
| Feeding difficulty | 133 (99.3%) | NA |
| Feeding tube used | 93 (69.4%) | NA |
| Weak cry | 131 (97.8%) | NA |
| Hypotonia | 132 (98.5%) | NA |
| Failure to thrive | 127 (94.8%) | 8.1%<sup>[9]</sup> |

PWS: Prader-Willi syndrome; BMI: body mass index; SD: standard deviation; NA: not available.

This cohort of patients had a higher rate of prematurity (16.4%) and low birth weight (32.8%) compared with 7.2% and 7.1%, respectively in China. The patients had a higher rate of birth asphyxia (32.1%) than the general population of neonates in 2014 as reported by Ministry of Health of China (6.3%) [6]. About 94.8% of the patients were hospitalized after birth with a median duration of 17 days (range: 0-90 days). 94.8% of the patients had failure to thrive which was far higher than general population (8.1%) [9]. Only 11.2% of the patients were purely breastfed at the first three months of life, and the rate is lower than the general population of 58.5% [9]. Feeding difficulty existed in 99.3% of the patients. 69.4% of the patients used feeding tube, however, 97.8% of them discontinued feeding tube and used silicone bottle, spoon even a syringe for feeding at home; and only two of them continued using feeding tube after hospitalization; 98.5% of the patients had hypotonia and 97.8% had weak cry even no cry at neonatal period.

All maternal and neonatal variables were compared between the patients with deletion and UPD (Table 2). Higher maternal age, maternal pre-pregnancy weight and BMI were noted in the UPD group than that in the deletion subgroup (all *P*<0.05). However, the rate of hospitalization at neonatal period and feeding tube used was higher in the deletion group than that in the UPD group with marginal differences (*P*=0.06 and *P*=0.11, respectively). We also compared our study with other large cohorts including two from France, one from UK and two from the United States (Table 3) [5,11–14]. Great similarities were found among these cohorts. Our cohort had a much higher rate of polyhydramnios and caesarean section compared with other cohorts.

**Table 2.** Comparison of the perinatal variables between the deletion and UPD groups

| Variables | Deletion (*n*=115) | UPD (*n*=19) | *P* value |
| --- | --- | --- | --- |
| **Maternal variables** |  |  |  |
| Maternal age (Year) | 29.6 ± 5.0 | 36.0 ± 6.1 | <0.001 |
| Maternal pre-pregnancy weight (kg) | 55.2 ± 8.3 | 59.9 ± 7.8 | 0.024 |
| Maternal weight gain during pregnancy (kg) | 12.1 ± 5.1 | 13.0 ± 3.7 | 0.44 |
| Maternal pre-pregnancy BMI (kg/m<sup>2</sup>) | 21.4 ± 3.0 | 23.3 ± 2.9 | 0.012 |
| Decreased fetal movements | 87.0% (94/108) | 93.7% (15/16) | 0.69 |
| Polyhydramnios | 42.6% | 42.1% | 0.79 |
| Oligohydramnios | 20.0% | 10.5% | 0.49 |
| Gestational diabetes | 5.9% | 0.0% | 0.31 |
| Gestational hypertension | 2.6% | 0.0% | 0.51 |
| Preeclampsia | 0.0% | 0.0% | - |
| Premature rupture of membranes | 10.4% | 5.3% | 0.69 |
| Caesarean section | 83.5% | 78.9% | 0.24 |
| **Neonatal variables** |  |  |  |
| Mean birth weight (≥37 weeks) (kg) | 2.82 ± 0.62 | 2.82 ± 0.43 | 0.99 |
| Mean birth weight (<37 weeks) (kg) | 2.40 ± 0.46 | 2.18 ± 0.71 | 0.48 |
| Low birth weight (<2.5 kg) ≥37 weeks | 25.2% | 21.0% | 1.00 |
| Birth length (cm) | 48.9 ± 3.4 | 48.9 ± 3.8 | 0.97 |
| Birth asphyxia | 34.0% | 21.0% | 0.30 |
| Hospitalization at neonatal period (Yes) | 96.5% | 84.2% | 0.06 |
| Breast feeding | 10.4% | 15.8% | 0.52 |
| Feeding difficulty | 99.1% | 100% | 0.81 |
| Feeding tube used | 72.2% | 52.6% | 0.11 |
| Weak cry | 98.3% | 94.7% | 0.37 |
| Hypotonia | 99.1% | 94.7% | 0.26 |
| Failure to thrive | 94.8% | 94.7% | 0.99 |

UPD: uniparental disomy; BMI: body mass index.

**Table 3.** Comparison with other reported large studies on perinatal variables in PWS

| Perinatal variables | China (N=134) | France (N=86)<br>(2006)<sup>[11]</sup> | United Kingdom<br>(N=46) (2008)<sup>[12]</sup> | France (N=61)<br>(2017)<sup>[5]</sup> | United States<br>(N=64) (2018)<sup>[13]</sup> | United States<br>(N=355) (2018)<sup>[14]</sup> |
| --- | --- | --- | --- | --- | --- | --- |
| Mean maternal age (year): Deletion | 29.6 | 29.3 | 31.4 | 31 | 28.7 | 29.2 |
| UPD | 36.0 | 36.4 | 37.9 | 38 | 36.7 | 35.2 |
| Decreased fetal movements | 109/124 (87.9%) | 47.6% | 67.4% | 27% | 85.6% | 78% |
| Polyhydramnios | 57 (42.6%) | 26.7% | 28.3% | 23.0% | NA | 18% |
| Vaginal delivery | 23 (17.2%) | 41.8% | 17.4% | 32.7% | 56.5% | 45.4% |
| Caesarean section | 111 (82.8%) | 53.4% | 52.2% | 67% | 42.1% | 54.6% |
| Preterm <37 weeks | 22 (16.4%) | 15% | 37% | 20% | 31.7% | 26% |
| Hypotonia | 132 (98.5%) | 96.5% | 100% | 76.1% | 84.3% | 99.7% |
| Feeding difficulty | 133 (99.3%) | 82.5% | 100% | 84.4% | 96.8% | 99% |

PWS: Prader-Willi syndrome; UPD: uniparental disomy; NA: not available.

## Discussion

PWS can now be diagnosed at a very early age or even at prenatal period benefiting from the improved molecular diagnosis techniques and increased awareness of PWS features [5,14–16]. Dobrescu et al and Gold et al reported that most patients still had a late diagnosis because the early signs of PWS not recognized, inappropriate molecular tests used, lacking of expertise in some hospitals and institutions [17,18]. Early diagnosis permits early care and treatment, which may reduce the hospital stay and tube feeding duration, hence preventing growth retardation and early onset of obesity [5,19–21]. To our knowledge, this is the first large cohort study on perinatal features of PWS patients in China.

Recognizing perinatal features of PWS will be helpful for early diagnosis and multidisciplinary care of affected infants. Our cohort had a high rate of polyhydramnios (42.5%), decreased fetal movements (87.9%), cesarean section (82.8%), low birth weight (32.8%), feeding difficulty (99.3%), hypotonia (98.5%), and failure to thrive (94.5%). The results were similar with previously reported cohorts from the UK, France and USA. Dudley et al reported high rates of polyhydramnios, induced labor and cesarean section, diminished fetal movement in a cohort of 86 French patients with PWS [11]. Gross et al found decreased fetal movements, small for gestational age, asymmetrical intrauterine growth and polyhydramnios in an Israel cohort [20]. Gold et al and Sign et al reported significantly higher rates of cesarean section, polyhydramnios, decreased fetal movements, low birth weight, feeding difficulty, hypotonia, and low Apgar scores in two cohorts of PWS patients in USA [13,14].

Hence, PWS should be highlighted for differential diagnosis for infants with following perinatal factors including polyhydramnios, intrauterine decreased fetal movements, cesarean section, low birth weight, feeding difficulty, hypotonia, and failure to thrive. Besides polyhydramnios, we also found a far higher rate of oligohydramnios (18.7%) in our cohort comparing with the general population (0.4-4.0%) in China. The accurate mechanism of polyhydramnios and oligohydramnios is still unclear although reduced fetal swallowing is regarded one cause of polyhydramnios; and high rate of oligohydramnios in this cohort maybe due to the high rate of premature rupture of memberanes. Further study for the mechanism of amniotic fluid disorder is required. Moreover, cesarean section rate was the highest in our cohort comparing with normal populations [7] and other cohorts [5,11–14]. The high caesarean section rate in this cohort was due to high rate of obstetric complications in PWS including abnormal amnion (61.2%), decreased fetal movements (87.9%), premature rupture of membranes (9.7%), or abnormal fetal heart rate/rhythm [21–23] although the data of fetal heart rate did not record in our study. High caesarean section rates may also be associated with the concepts and selection of pregnant women in China.

Considering neonatal complications, failure to thrive, low birth weight, feeding difficulty and hypotonia were most common in our study. Moreover, rate of asphyxia was higher in our cohort than that in general population in China (32.1% vs. 6.3%) [6]. This may be associated with higher ratio of intrauterine abnormal fetal heart rate/rhythm and decreased fetal movements, hypotonia and weak cry after birth that may be considered as intrauterine or intrapartum asphyxia, and higher ratio of preterm in our cohort as well. More than 99% of the infants had feeding difficulty requiring tube feeding and about 70% of the infants ever used feeding tube during hospitalization. Tube feeding was discontinued in 98% of these infants by their parents. Parents refused to use tube feeding at home; they preferred using silicone bottle, spoon even a syringe for feeding PWS infants which cannot reach the effect of tube feeding. As a result, the rate of failure to thrive (94.5%) during the first months of life in this cohort of patients was higher than that reported by Sign et al (77%) [14]. The possible reasons for refusing tube-feeding at home may be lacking education of tube-feeding from healthcare workers and parents’ not familiar with home tube feeding techniques. Therefore, it is very important to strengthen feeding knowledge and ability of parents of PWS patients in China.

Compared the variables between the deletion and UPD subgroups are still conflicting in literature. Gillessen-Kaesbach et al and Whittington et al showed that UPD patients had a significantly higher birth weight and maternal age than that of deletion patients [12,23]; vise versa, Gunay-Aygun et al found a significantly lower birth weights and lengths in the UPD group than those in the deletion group [23]. Dudley and Muscatelli found a higher rate of induced labor, premature labor and higher maternal age in the UPD group, and a high rate of low birth weight in the deletion group [11]. However, Sign et al only find higher maternal age and pre-pregnancy weight in the UPD group in a large cohort of 355 PWS patients from USA [14]. Whittington et al found significant difference of maternal age and birth weight between different genetic subtypes [12]. In our study, higher maternal age, pre-pregnancy weight and BMI were noted in the UPD subgroup that that in deletion subgroup; however, the rate of hospitalization at neonatal period and feeding tube used was higher in the deletion group with marginal differences (0.05<P<0.15). These implied that patients with deletion genetic type may have more difficulty in feeding, and prone to be hospitalized at neonatal period. Large cohort study is required to confirm these findings.

The mechanism of PWS is still unclear. We found that high maternal age of PWS patients, especially in the UPD group, which was reported in several reports [12, 14, 24], which implying that the advanced maternal age is associated with PWS resulting from non-disjunction at meiosis 1. Moreover, we also noted that mothers of PWS patients had higher pre-pregnancy BMI than those in the general population, and higher pre-pregnancy weight and BMI in UPD group with marginal differences, which was also reported in a large cohort of 355 PWS patients from USA [14]. These suggested that high maternal age and overweight may be the risk factors of PWS and further studies for the mechanism of PWS are required.

In conclusion, our study highlighted the perinatal features of a large cohort of Chinese patients with PWS. We found high incidence of decreased fetal movements, polyhydramnios and delivery by cesarean section, and higher maternal age comparing with the general population. Neonatal features found in our cohort included low birth weight, birth asphyxia, failure to thrive, feeding difficulty, weak cry and hypotonia. We found mothers of patients with UPD had higher maternal age, pre-pregnancy weight and BMI as comparing with the deletion group. Most patients required tube feeding during hospitalization, however tube feeding was discontinued by their parents after discharge at home. Nutrition deficiency was a serious problem in infants with PWS. Home tube-feeding instruction should be carried out for parents of PWS patients in China. Our study helps to better understanding of the perinatal features of PWS in China, which will benefit for early diagnosis and treatment. And our results provide some evidence for further including PWS screening into the newborn screening program in China.

## Methods

### Subjects

This study is part of a project started by PWS Research Group from Children’s Hospital, Zhejiang University School of Medicine. The PWS Research Group is establishing a PWS Registry for patients in China. The study was approved by Institutional Review Board of Zhejiang University School of Medicine (no. 2018-IRB-055).

Written informed consents were obtained from all parents or the patients (above 7 years) registered in the PWS Registry. We analyzed anonymous data of 134 patients from the PWS Registry. Perinatal and neonatal presentations were analyzed, and compared between the two PWS genetic subtypes as well as general population [6–10] and data reported in other countries [5,11–14].

### Statistical analysis

All data were analyzed in the SPSS (version 16.0) software package. The categorical variables were summarized using frequencies, and continuous variables calculated mean ± SD scores. Student’s *t*-test was used to compare continuous variables, and Pearson *χ^2^* test was used for comparing categorical variables. Fisher’s exact test was used to compare categorical variables when the expected cell count was less than 5. A two-tailed *P* value less than 0.05 was considered statistically significant.

## Acknowledgements

We thank all parents and children who were registered in the PWS registry for supporting our research.

## Author Contributions

YL, ZQ, and ZC designed the study; YL, ZQ collected the data; YL, ZQ, MB, MS, DY and ZM analyzed and interpreted data. YL wrote the first draft of the paper. ZQ, MB and ZC revised the paper critically. YL and ZQ contributed equally to this paper. All authors have participated in revising the manuscript critically and gave their final approval of the version to be submitted.

